# Direct estimate of the spontaneous mutation rate uncovers the effects of drift and recombination in the *Chlamydomonas reinhardtii* plastid genome

**DOI:** 10.1101/031898

**Authors:** Rob W. Ness, Susanne A. Kraemer, Nick Colegrave, Peter D. Keightley

## Abstract

Plastids perform crucial cellular functions, including photosynthesis, across a wide variety of eukaryotes. Since endosymbiosis, plastids have maintained independent genomes that now display a wide diversity of gene content, genome structure, gene regulation mechanisms, and transmission modes. The evolution of plastid genomes depends on an input of *de novo* mutation, but our knowledge of mutation in the plastid is limited to indirect inference from patterns of DNA divergence between species. Here, we use a mutation accumulation experiment, where selection acting on mutations is rendered ineffective, combined with whole-plastid genome sequencing to directly characterize *de novo* mutation in *Chlamydomonas reinhardtii*. We show that the mutation rates of the plastid and nuclear genomes are similar, but that the base spectra of mutations differ significantly. We integrate our measure of the mutation rate with a population genomic dataset of 20 individuals, and show that the plastid genome is subject to substantially stronger genetic drift than the nuclear genome. We also show that high levels of linkage disequilibrium in the plastid genome are not due to restricted recombination, but are instead a consequence of increased genetic drift. One likely explanation for increased drift in the plastid genome is that there are stronger effects of genetic hitchhiking. The presence of recombination in the plastid is consistent with laboratory studies in *C. reinhardtii* and demonstrates that although the plastid genome is thought to be uniparentally inherited, it recombines in nature at a rate similar to the nuclear genome.

## Introduction

Plastids, including the chloroplast, arose via the primary endosymbiosis of a cyanobacterium and subsequently spread laterally into other lineages through eukaryote-eukaryote endosymbioses. The genome of the original endosymbiont has been retained since the the ancestor of glaucophytes, red algae and green plants (but see Molina et al. 2014; Smith and Lee 2014). The near ubiquity, sequence conservation and high copy number of plastid DNA has made it an invaluable resource for DNA barcoding and biogeographical/phylogenetic investigations. Most of our knowledge of plastid biology and genomics is derived from a disproportionate sampling of land plant chloroplasts, but recent sampling of a broader array of taxa has revealed an extraordinary diversity of genomic architectures and modes of inheritance (reviewed in Smith and Keeling 2015). The structure and composition of plastid genomes often differ significantly from their nuclear and mitochondrial counterparts. Despite the widespread use of plastid genomes in evolutionary genetics, we know little about the forces that explain diversity and influence their molecular evolution.

Variation in the type and rate of mutation among the nuclear, mitochondrial and plastid genomes has been posited as the source of differences in their genome structure and evolution (Allen and Raven 1996; Race et al. 1999; Lynch et al. 2006). However, despite a growing body of information on spontaneous mutation in the nucleus (eg. Kong et al. 2012; Zhu et al. 2014) and mitochondria (e.g. Haag-Liautard et al. 2008; Lynch et al. 2008; Howe et al. 2010) we know relatively little about spontaneous mutation in plastids. Most, if not all, DNA repair and replication genes used in the plastid are nuclear-encoded, some of which are dually targeted (e.g. *PHR1;* Petersen and Small 2001), which suggests that patterns of mutation might be similar. However, biochemical processes in the plastid and mitochondria create mutagenic reactive oxygen species and may lead to a fundamentally different mutational environment for organellar genomes. Mutagenesis by oxidative stress has been implicated in cellular aging (Van Breusegem and Dat 2006), the transfer of plastid genes to the nucleus (Allen and Raven 1996), and the retention of particular proteins in the organelle genome to maintain redox balance (Race et al. 1999). Moreover, the plastid and mitochondrial genomes replicate multiple times per cell division and this could increase the mutation rate if mutations were introduced during DNA replication.

Virtually all of our knowledge of mutational processes in plastids is derived from patterns of DNA substitution between species. In a strictly neutral stretch of DNA, the amount of divergence between a pair of species is proportional to the mutation rate and the time since divergence (Kimura 1983). Analysis of organelle genome divergence has been a fruitful approach and yielded some general conclusions about mutation in plastid genomes. In land plants, most evidence suggests that plastid genomes have a lower mutation rate than their nuclear counterpart, but higher than the mitochondrial genome (for review see Smith and Keeling 2015). The lower plastid mutation rate may be limited to land plants, however, and the opposite appears to be true in most other taxonomic groups (Smith 2015). For example, in red algae and glaucophytes, putatively neutral substitution rates in the plastid are similar to the nuclear genome, but much lower than the rate of substitution in the mitochondria. In green algae, the focus of this study, divergence in the plastid, mitochondrial and nuclear genomes is remarkably similar (Smith and Keeling 2012).

There are a number of difficulties with using between-species nucleotide divergence for estimating mutation rates (for a review see Sloan and Taylor 2011). Primarily, the assumption that nucleotide substitutions are neutral is unlikely to be true. Synonymous sites are frequently used as a proxy for neutral sites, and sometimes combined with intronic or intergenic sites (often referred to as ‘silent’ sites). However, there is clear evidence that synonymous sites are under selection related to codon usage bias, and RNA secondary structure (Chamary et al. 2006). Silent sites may also be influenced by non-neutral processes, such as GC-biased gene conversion that are heterogeneous across the genome(s) and between species, complicating the interpretation of substitution rates (Lesecque et al. 2013). In addition, the base composition of synonymous sites is distinct from the genome background, and, given that the mutation rate of the four bases is variable and dependent on the nucleotide sequence surrounding a given site, synonymous sites are expected to have their own mutational properties that may vary between nuclear and organellar genomes (Ness et al. 2015). Lastly, phylogenetic-based estimates of substitution need to account for multiple mutations occurring at the same site, which is especially problematic when comparing distantly related taxa. Phylogenetic methods are also prone to violations of their assumptions, such as the presence of recombination, lateral gene transfer and intraspecific polymorphism (see Lanfear et al. 2010; Sloan and Taylor 2011). More direct measurements of the properties of spontaneous mutation are therefore needed to assess how well patterns of substitution reflect the process of mutation in plastid genomes.

One method to circumvent the complications of inferring mutation from substitution and to disentangle mutation rate from divergence time is to measure *de novo* mutations directly in mutation accumulation (MA) experiments. In a MA experiment populations are maintained for many generations under minimal natural selection to allow mutations to accumulate regardless of their fitness consequences. Increasing the strength of genetic drift by regularly bottlenecking the population allows random, unbiased accumulation of all but the most strongly deleterious mutations. When MA is combined with genome sequencing, we can directly study of the process of mutation (eg. Zhu et al. 2014; Ness et al. 2015). Data from direct estimates of mutation rate in mitochondria suggest that substitution-based estimates of mutation may be as much as two orders of magnitude lower than the true rate (Denver et al. 2000; Howell et al. 2003; Haag-Liautard et al. 2008; Lynch et al. 2008). Data of this kind do not exist for plastid genomes because, at least in land plants, they are thought to have lower mutation rates than mitochondria and therefore require many generations of MA and/or sequencing of many mutated genomes to observe *de novo* mutations. Two previous mutation accumulation experiments that estimate the mutation rate in *C. reinhardtii* failed to detect any mutations in the plastid genome (Ness et al. 2012; Sung et al. 2012).

Estimates of the spontaneous mutation rate can be combined with patterns of genetic variation to describe key properties of species’ history and population genetics. Following Kimura (1983), genetic divergence between two species is twice the product of the mutation rate and the number of generations (*t*) since speciation (*2μt*). A direct estimate of mutation rate can therefore be used to calculate divergence times between species, and identify regions of the genome where selection has increased or decreased the rate of molecular evolution. In addition, the amount of neutral genetic variation maintained in a population (*θ*) is determined by the strength of drift, measured as the effective population size (*N_e_*) and the mutation rate via the relationship *θ* = *2N_e_μ* (*4N_e_μ* in diploids). We can therefore use the mutation rate to infer *N_e_* from natural levels of polymorphism. The effective population size is a fundamentally important property of a population that has a strong impact on the efficacy of selection (2*N_e_s*) and the breakdown of linkage disequilibrium (LD) (population recombination rate, *ρ* = 2*N_e_r*, where *r* is the rate of recombination per base per generation). Thus, the integration of laboratory-based estimates of mutation with population genomic sequencing from natural samples can facilitate investigation into the biology of species not possible with either approach in isolation.

In this study, we present the first direct estimate of the spontaneous mutation rate in a plastid genome by combining an MA experiment with sequencing of the plastid genome of *Chlamydomonas reinhardtii*. We combine our estimate of mutation rate with whole-plastid sequencing of 20 natural isolates of *C. reinhardtii* to investigate patterns of drift and recombination. *Chlamydomonas reinhardtii* is a unicellular green algae with a plastid genome of 204kbp. It can reproduce both sexually and asexually. Sex occurs by fusion of opposite mating types (mt+ and mt-), after which the plastid from the mt-parent and the mitochondria from the mt+ parent are eliminated, resulting in a primarily uniparental mode of organellar inheritance. Comparisons of divergence between *C. reinhardtii* and *C. incerta* genomes suggest that, unlike land plants, the plastid mutation rate of *C. reinhardtii* is similar to the rate in the nuclear and mitochondrial genomes (Hua et al. 2012). However, for the above mentioned reasons it is difficult to know whether the patterns of substitution reflect real differences in the mutation between the three genomes. In this paper we address three questions (1) What is the mutation rate of the plastid genome of *C. reinhardtii?* (2) How does the rate and spectrum of plastid mutation compare with the nuclear genome? (3) Using the mutation rate, what is the effective population size of the *C. reinhardtii* plastid and nuclear genomes and how does this relate to recombination?

## Materials and Methods

**Mutation accumulation experiment**. We conducted a mutation accumulation experiment in six genetically diverse strains of *C. reinhardtii* obtained from the Chlamydomonas Resource Center (chlamycollection.org). The strains were chosen to broadly cover the geographic range of known *C. reinhardtii* samples in North America (Table S1). To initiate the MA lines, a single colony from each of the six ancestral strains was streaked out, and we randomly selected 15 individual colonies to start the replicated MA lines (90 MA lines total). We bottlenecked the MA lines at regular intervals by selecting a random colony that was streaked onto a fresh agar plate. We estimated the number of generations undergone by each MA line over the course of the experiment by measuring the number of cells in colonies grown on agar plates after a period of growth equivalent to the times between transfers in the experiment. Characterization of these MA lines was published in Morgan et al. (2014) and whole genome sequencing and characterization of the nuclear mutation rate was published in Ness et al. (2015).

**Population genomic sequencing**. Twenty natural isolates of *C. reinhardtii* sampled from two fields approximately 75km apart in Quebec, Canada, were acquired from the *Chlamydomonas* collection (www.chlamycollection.org). These *C. reinhardtii* strains are the only collection where more than two individuals from a single locality were collected. A within-location sampling strategy allowed better estimation of population genetic parameters, such as the effective rate recombination (*ρ=2N_e_r*) and genetic diversity (*θ_π_=*2*N_e_μ*), than a species-wide sample, which may be confounded by the effects of population structure (Städler et al. 2009).

**Sequencing and alignment**. DNA was extracted from frozen, concentrated cultures using a standard phenol-chloroform extraction. We conducted whole-genome re-sequencing on the Illumina GAII platform at the Beijing Genomics Institute (BGI-HongKong Co., Ltd, Hong Kong). We modified the library amplification to accommodate the unusually high GC content of the *C. reinhardtii* genome (mean GC= 63.9%) following (Aird et al. 2011) (3 min at 98°C; 10 × [80 sec at 98°C, 30 sec at 65°C, 30 sec at 72°C]; 10 min at 72°C and slow temperature ramping 2.2°C/sec). We obtained an average of 28.6× coverage of each sampled genome (3Gbp of 100bp paired-end sequence) and because the plastid is present at high copy number in the cell, mean coverage of the plastid was 1034×.

We aligned sequencing reads using BWA 0.7.4-r385 (Li and Durbin 2009) to the *C. reinhardtii* reference genome (version 5.3). Because the reference genome does not include the organelles, we added the plastid genome (NCBI accession NC_005353), mitochondrial genome (NCBI accession NC_001638) and the MT-locus (NCBI accession GU814015). To call genotypes, we used the GATK v3.3 (Depristo et al. 2011) tool ‘HaplotypeCaller’ which performs local re-assembly of reads around problematic areas to more accurately infer single nucleotide variants and short indels (<50bp). All replicated MA lines derived from a given ancestral strain and the 20 natural lines from Quebec were run in HaplotypeCaller jointly. The resulting Variant Call Format files (VCFs) (one per ancestral strain and one for the natural lines) were converted to Wormtable databases using Wormtable v0.1.0 (Kelleher et al. 2013) to enable efficient mutation identification and population genetic analysis.

**Mutation rate estimation**. Replicate MA lines derived from a given ancestral strain were initially genetically identical, therefore unique alleles carried by a single replicate were considered candidate mutations. We applied the following filters to candidate mutations identified in the plastid genome to minimize erroneous calls:

1. Mapping quality (MQ) ≥ 90 and PHRED scaled site quality (QUAL) ≥ 100
2. All lines called ‘homozygous’; *C. reinhardtii* plastid is haploid and this filter eliminates mapping errors from paralogous loci.
3. The genotype of exactly one MA line differed from the rest of the lines
4. All non-mutated lines shared the same genotype
5. At least two MA lines have genotype calls

**Heteroplasmy**. We investigated the possibility that spontaneous mutations may be heteroplasmic within an MA line. Alleles at heteroplasmic sites are not expected to segregate at 1:1, like heterozygote sites in diploids, and will not be accurately called by standard variant calling algorithms. We therefore attempted to detect heteroplasmic sites directly from raw read data, extracted using SamTools mpileup (Li et al. 2009). We applied the following filters to each group of replicate MA lines from a common ancestor:

1. One individual has ≥50x coverage, ≥5% of which carry a minor allele
2. At least two MA lines had reads aligned to the position
3. Other samples have <2 aligned reads that carry the candidate mutant allele

We estimated the accuracy of our mutation calls using Sanger sequencing of all candidate mutations including both indels and single nucleotide mutations (SNMs). We amplified each locus in the putative mutant MA line and a non-mutated line from the same ancestral strain. Sequences were then visually inspected in SeqTrace v0.9.0 to confirm the presence of the mutated site.

We calculated the mutation rate as, *μ* = mutations / (high quality sites × MA generations). To obtain accurate mutation rate estimates, it is necessary to define the number of sites at which a mutation could have been detected. Such ‘callable’ sites were determined using the same criteria outlined above except we required all MA lines from a given ancestor share the same genotype and the minimum PHRED-scaled QUAL score was 36 to account for the difference in QUAL distributions between invariant and variant sites (see Ness et al. 2015). To analyze mutation spectrum, we treated complementary mutations (C:G and A:T) symmetrically, such that there were six distinct SNMs (A:T→C:G, A:T→G:C, A:T→T:A, C:G→A:T, C:G→G:C, C:G→T:A). To assess the base spectrum of mutation, we calculated the frequency of each mutation type relative to a random expectation adjusted for the base composition of the callable sites in the plastid genome. We also compared the plastid and nuclear mutation spectrum reported by Ness et al. (2015).

**Population genetic analysis**. We estimated nucleotide diversity (*θ_π_* = 2*N_e_μ*) in the plastid and nuclear genomes of the 20 natural lines from Quebec. We estimated neutral diversity using silent sites (4-fold degenerate, intronic and intergenic sites) and we estimated selective constraint as the ratio of diversity at 0-fold and 4-fold degenerate sites (*θ*_0_ / *θ*_4_). We inferred the effective population size (*N_e_*) for the plastid and nuclear genomes using their respective levels of silent diversity (*θ_π_*) and mutation rates (*μ*) as *N_e_=θ_π_/*2*μ*. If the plastid genome is uniparentally inherited, all else being equal, it is expected to have approximately one-half the *N_e_* of the nuclear genome. Using 4-fold degenerate sites to estimate "neutral” diversity suffers from the problems explained in our introduction, therefore our estimates of neutral diversity may be underestimated. To test whether the *θ_π_* for nuclear and plastid genomes differ, we calculated *θ_π_* for sliding windows of 10kbp in both genomes. We then bootstrapped the sliding windows 1,000 times with replacement and for each bootstrap replicate we calculated the difference in mean *θ_π_* between the plastid and nuclear genomes. The proportion of bootstrap replicates with differences in mean *θ_π_* that overlap zero indicate whether the plastid and nuclear genomes have significantly different mean levels of nucleotide diversity.

We inferred variation in the effective rate of recombination (*ρ* = 2*N_e_r*) from polymorphism data with the program LDhelmet (Chan et al. 2012). LDhelmet uses a similar reverse-jump Markov Chain-Monte Carlo approach as employed in LDHat (Mcvean et al. 2004), but includes modifications to improve recombination rate estimates and computational efficiency, especially when there are missing data. We generated haplotype configurations with the recommended window size of 50 SNPs (-w 50). To calculate the likelihood look-up table and Padé coefficients, we used the mean 4-fold degenerate diversity measures from the nuclear and plastid genomes for the *θ* parameters (-t 0.039 and 0.003, respectively). To estimate whether *ρ* varied between the nuclear and plastid genomes we calculated *ρ* per base in sliding windows of 10kbp. We then used the same bootstrapping method described above for nucleotide diversity to determine whether there was a significant difference between *ρ* in the plastid and nuclear genomes.

We estimated *N_e_* as *θ_π_/*2*μ* and the recombination rate *r* as *ρ*/2*N_e_*. We approximated error in our *N_e_* estimate by resampling both the mutation rate and *θ_π_*. For mutation rate, we drew 1,000 samples from a Poisson distribution with a mean equal to the number of observed mutations in the nuclear and plastid genomes. We then calculated *μ* for each draw by dividing it by the number of sites and generations. We then estimated error in *N_e_* by dividing the mean of each of 1,000 bootstrap samples of *θ_π_* by twice each sampled mutation rate. The standard error was calculated as the standard deviation of the 1,000 samples. We then used the 1,000 *N_e_* values to estimate the standard error of the recombination rate (*r*) by dividing each of the 1,000 bootstrap samples of *ρ* by twice each sampled value of *N_e_*.

## Results & Discussion

**Plastid Mutation rate**. In total, across all 85 MA lines, we analyzed 1.38 × 10^7^ plastid genome sites over an average of 929 generations (mean coverage of each line = 1,565, from 317 to 4,743). On average, 163kbp of the 204kbp plastid genome was callable in each line and the missing sites were accounted for by the two ~19kbp inverted repeats, where ambiguous alignments made genotypes difficult to call (Figure 1). We identified 12 *de novo* mutations, including 10 SNMs and two short insertions (Figure 1, Table 1). We confirmed 9 of the 10 SNMs and the two insertions using Sanger sequencing. The unconfirmed SNM appeared very well aligned in the Illumina read data (see supplementary figure 1), but the region in which it occurs is flanked by repetitive DNA that made PCR amplification of a single locus difficult. We therefore treated this SNM as a true mutation, yielding a total mutation rate of *μ*_total_ = 9.23 × 10^−10^ (Poisson 95% CI 4.8-16.1 × 10^−10^) with an SNM rate *μ*_SNM_ = 7.69 × 10^−10^ (excluding the unconfirmed SNM yields, *μ*_total_ = 8.46 × 10^−10^ and *μ*_SNM_ = 6.92 × 10^−10^).

**Figure 1.**
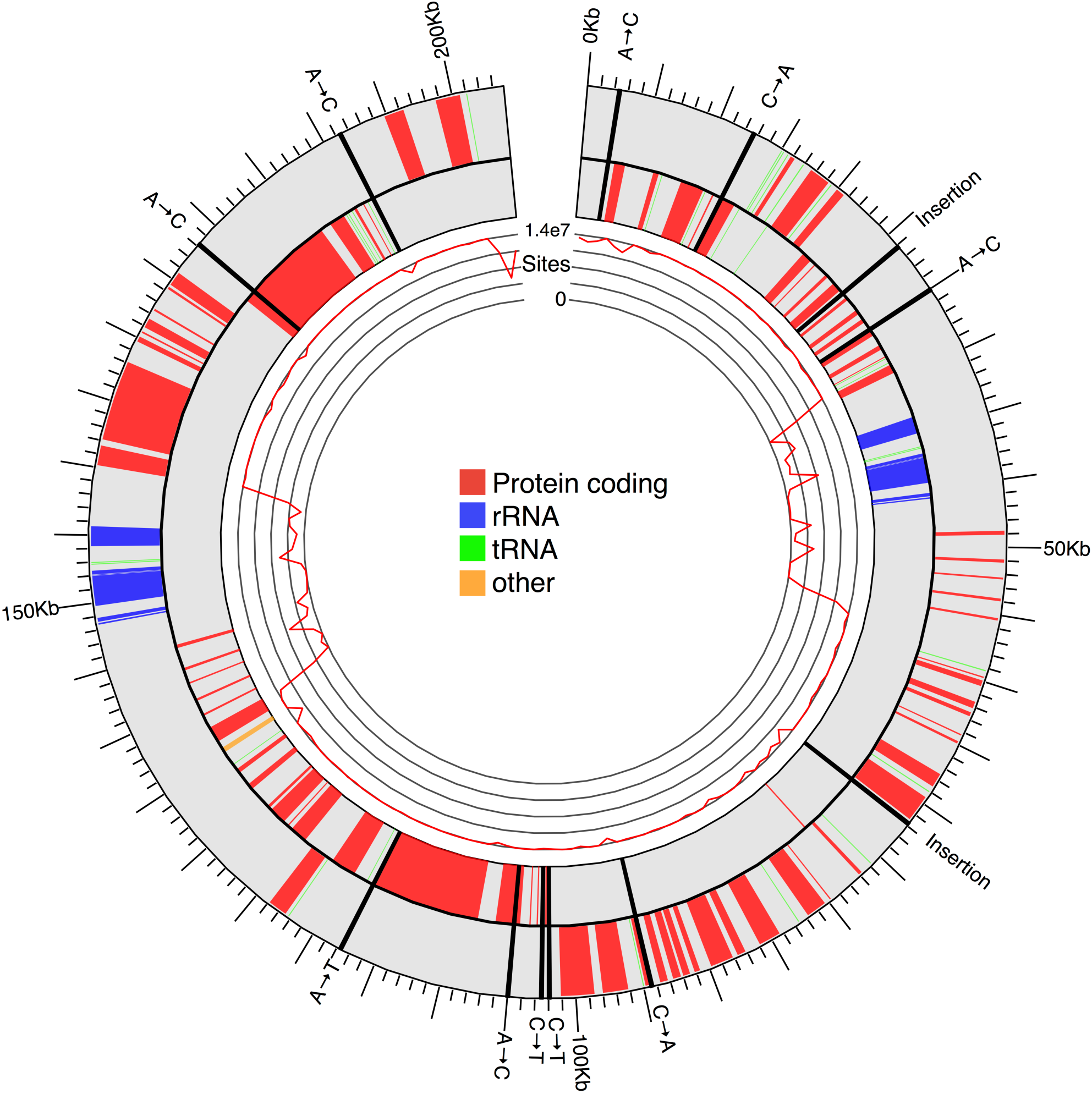
Ideogram of the *Chlamydomonas reinhardtii* plastid genome showing locations of *de novo* mutations. The shaded grey circle represents the 204Kb plastid genome, showing protein-coding (red), tRNA (green) and rRNA (blue) exons as coloured boxes. Genes on the positive strand are shown on the inner track and genes on the negative strand on the outer track. Mutations are marked as thick black lines across the ideogram and labelled with the mutation event. The inner circle shows the total number of high quality plastid sites sequenced multiplied by the total number of generations of mutation accumulation experienced in all 85 lines. The dips in the number of high quality sites between 35–55Kb and 138–158Kb correspond to the two copies of the inverted repeat (IR) region.

**Table 1.**
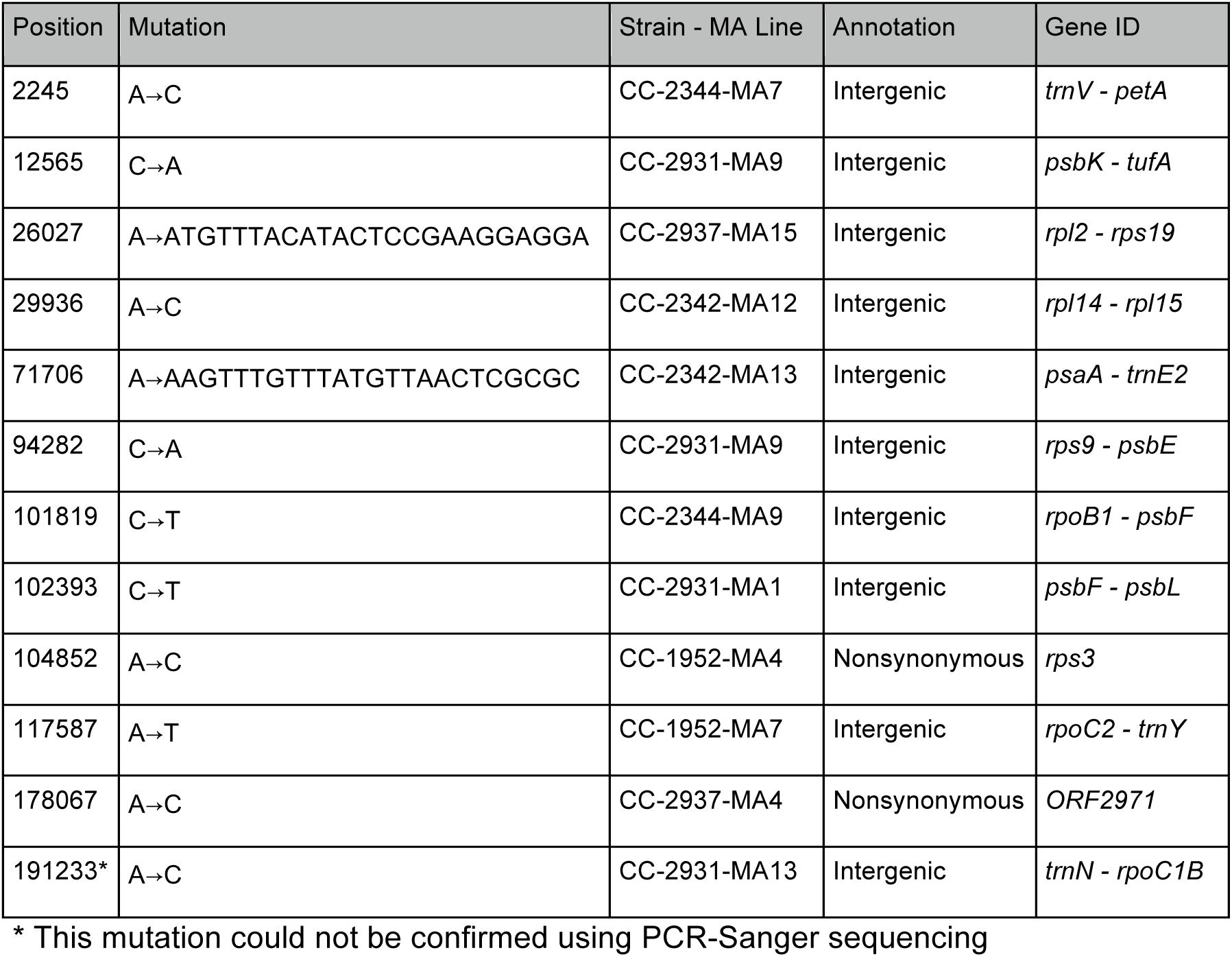
Table of de novo mutations identified in the plastids of 85 MA lines of *Chlamydomonas reinhardtii*. The strand of each mutation was converted so that ancestral state is an A or C. For intergenic mutations, the two genes flanking each mutated site are indicated, and for the nonsynonymous mutations the affected gene is reported.

Both the total and SNM plastid genome mutation rates are marginally lower, but not significantly different from, the nuclear genome mutation rate (*μ*_nuclear total_ = 11.5 × 10^−10^, *μ*_nuclear SNM_ = 9.63 × 10^−10^), estimated in the same sets of 85 MA lines (Ness et al. 2015). It is also worth noting that we did not observe any mutations in the mitochondrial genome. If the much smaller mitochondrial genome (15.6 kbp) had the same mutation rate as the plastid genome, we would expect to observe only 1.15 mutations. Treating the observed number of mutations as a Poisson variable, we calculated the upper limit on the mitochondrial mutation rate to be *μ*_mtDNA_ < 3.0 × 10^−9^. We also investigated our data for evidence of heteroplasmy. To calculate the number of heteroplasmic mutations that we would expect to detect, we considered an individual MA line as a population of plastid genomes in which we can calculate the probability of observing a segregating mutation from the predicted diversity within a population, 2*N_e_μ*. In this case, *N_e_* is the effective number of plastid genome copies transmitted through each vegetative cell division, approximately 10 (5-15; Vanwinkle-Swift 1980; Birky 2001). Across all 85 MA lines we expect to observe 0.26 heteroplasmic mutations (2 × *N_e_* × *μ* × total callable sites). Not surprisingly, given the small number of mutations detected, we found no evidence of heteroplasmic mutations across the 85 MA lines.

The overall similarity of the mutation rate between the nuclear and plastid genomes is consistent with similar levels of synonymous nucleotide divergence in the nuclear and plastid genomes between *C. reinhardtii* and *C. incerta* (Smith 2015). Thus, even if selection or other forces, such as biased gene conversion, influence silent site divergence, their effects are similar in the nuclear and plastid genomes. Unfortunately, we cannot estimate the extent to which the rate of synonymous substitution is reduced by selection because no independent estimate of the divergence time between *C. reinhardtii* and *C. incerta* exists. The similarity of plastid and nuclear mutation rates in *C. reinhardtii* is strikingly different from most land plants, where plastid substitution rates are substantially lower than nuclear substitution rates (Smith and Keeling 2015). However, without direct characterisation of mutation in the plastids of land plants, we cannot know whether the difference in substitution rates reflects differences in mutation rates or differences in the strength of selection on silent sites.

**Base spectrum of plastid mutation**. There is a notable difference in base spectrum of mutations between the plastid and nuclear genomes. Although the total number of mutations was small, the base spectrum of the plastid is significantly different from the nuclear genome (*χ*^2^(5,10) = 11.4, *p* < 0.05).

While the nuclear genome is strongly biased towards C:G → T:A transitions, in contrast the plastid genome SNMs show an excess of A:T → C:G and C:G → A:T mutations and a deficit of A:T → G:C (from Ness et al. 2015). The excess of A:T → C:G transversions is consistent with a study that examined mutation in three positions of an Aat II restriction site in the *C. reinhardtii* plastid 16S sequence (GuhaMajumdar and Sears 2005). Using the population genomic data, we also examined the composition of naturally occurring polymorphisms in the plastid to assess whether their base spectrum is consistent with our inference from *de novo* mutations. Specifically, low frequency polymorphisms at 4-fold degenerate sites will have had less time to be influenced by selection, and should therefore reflect the base spectrum of mutations. In accordance with the *de novo* mutations there is an excess of A:T → C:G singleton and doubleton polymorphisms at 4-fold degenerate sites (observed/expected = 1.5), but there is an even stronger bias towards mutations at C:G sites (observed/expected = 2.8). Both the base composition of *de novo* mutations and 4-fold singletons predict that the equilibrium GC content of the plastid would be 39.7% which is close to the observed value of 34.5%. This contrasts with the nuclear mutation spectrum, which would predict an equilibrium GC content of 29% (Ness et al. 2015). Interestingly, the base composition of 4-fold degenerate sites in the plastid genome is 89.2% AT, suggesting strong selection on codon usage in plastid genes.

**Integrating mutation rate estimates with population genetics**. We can use our estimate of the mutation rate to quantify the time since divergence from other species. If substitutions are neutral, then the expected divergence between two species is *2μ*t, where *t* is the time in generations since the split. Based on a divergence value of 0.30 for synonymous sites in the plastid (Hua et al. 2012) the split between *C. reinhardtii* and *C. incerta* therefore occurred ~1.93 × 10^8^ generations ago. Unfortunately, we know little to nothing about the number of generations per year in nature and therefore cannot speculate on the amount of time that separates these species. As a result, it is not possible to ascertain the extent to which estimates of neutral substitution rate and spontaneous mutation rate differ.

Our estimate of the mutation rate can reveal other aspects of the population biology and evolutionary history of *C. reinhardtii*. Equilibrium neutral nucleotide diversity (*θ_π_*) is proportional to product of the mutation rate and effective population size (*θ* = *2N_e_μ*). The effective population size reflects the amount of genetic drift in a population, specifically the number of individuals in an idealized Wright-Fisher population that would display the observed amount of drift. *N_e_* can therefore be altered by population size fluctuations, bottlenecks and natural selection. Among the 20 natural isolates from Quebec, plastid genome neutral genetic diversity was seven-fold lower than that of the nuclear genome (plastid *θ_π_* = 0.0039, nuclear *θ_π_* = 0.0275, bootstrap test *p* < 0.001; Figure 2). If the plastid genome is uniparentally inherited, all else being equal, it is expected to have approximately one-half the *N_e_* of the nuclear genome. The SNM mutation rate estimates from the plastid (7.8 × 10^−10^) and nuclear genomes (9.65 × 10^−10^; Ness et al. 2015) imply that *N_e_* for the plastid genome is 2.8× lower than the predicted two-fold reduction (plastid *N_e_* = 2.5 × 10^6^, SE 1.6 × 10^6^, nuclear *N_e_* = 1.4 × 10^7^ SE 2.0 × 10^5^, bootstrap test *p* < 0.01, Figure 2). Given that the plastid genome is thought to be largely non-recombining, hitchhiking, i.e., by selection for advantageous alleles (sweeps) or selection against deleterious alleles (background selection), may reduce plastid *N_e_* more severely than the nuclear genome. Moreover, strong effects of hitchhiking could be observed in the presence of recombination if the plastid is subject to more frequent and stronger selection than comparable regions of the nuclear genome.

**Figure 2.**
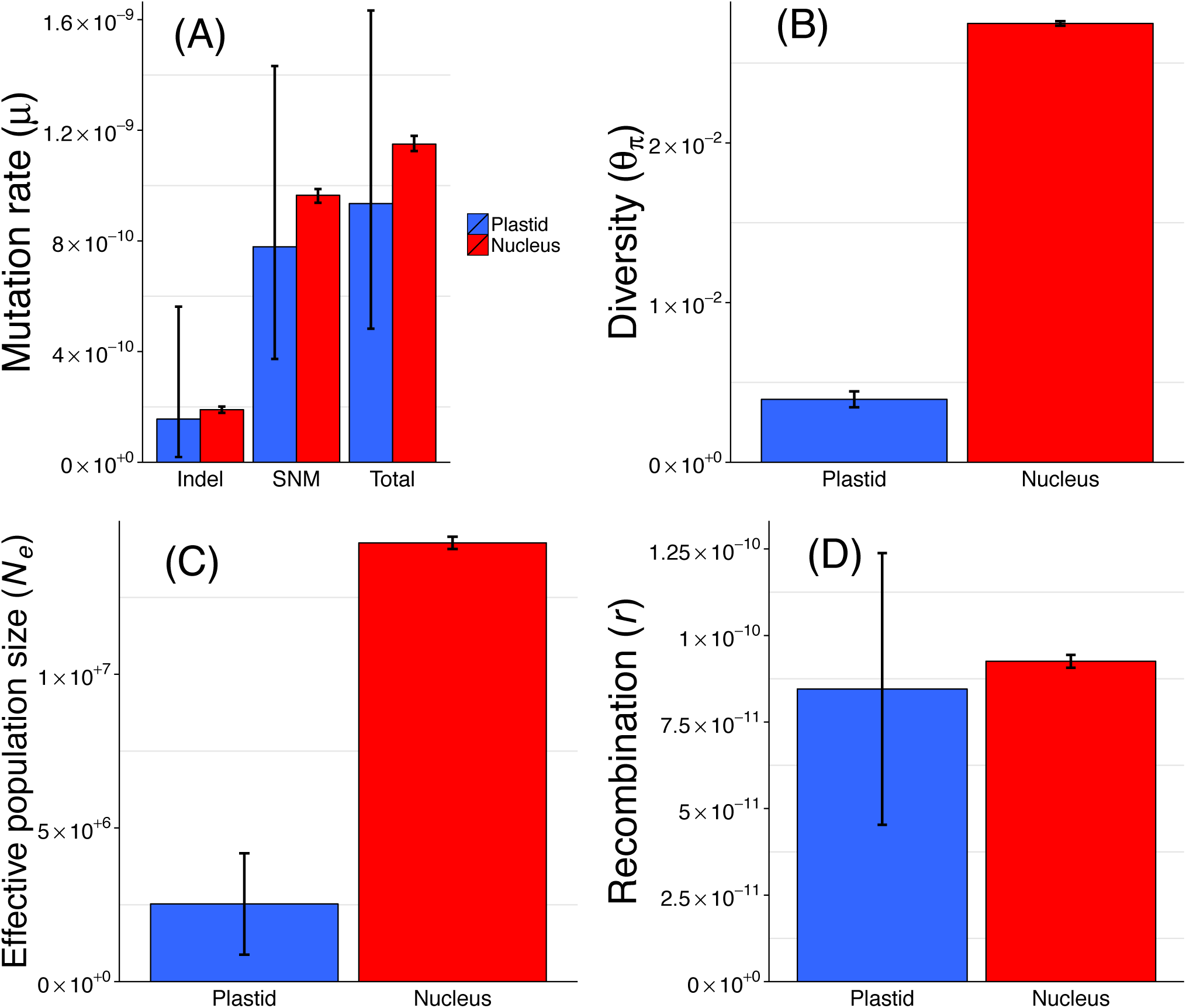
Comparison of nuclear and plastid (A) mutation rate, (B) nucleotide diversity, (C) effective population size (N_e_) and (D) recombination rate (r). Nuclear mutation rates were taken from Ness et al. (2015). *N_e_* was estimated using diversity (*θ_π_*) estimates and SNM mutation rates with the formula *N_e_=θ_π_/*2*μ*. The estimated plastid and nuclear *N_e_* values were then used to calculate recombination rates per base pair per generation (r) from the equation *r* = *ρ/*2*N_e_*, where *ρ* is the effective recombination rate estimated from patterns of linkage disequilibrium in polymorphism data using the program LDHelmet (Chan et al. 2012). Error bars on the mutation rate estimates are 95% confidence intervals assuming mutation was a Poisson process. Error bars on panels (B), (C), and (D) represent ±1 standard error, estimated using a bootstrap procedure. Error bars are larger for the plastid compared to the nucleus because there are fewer sites and mutations in the plastid genome.

Having an estimate of *N_e_* based on the mutation rate and genetic diversity, we can use patterns of linkage disequilibrium (LD) to test the assumption that the rate of recombination is indeed lower in the plastid. During sexual reproduction in *C. reinhardtii*, there is a period after the gametes fuse, but before the mt-plastid is eliminated, that provides an opportunity for recombination to occur. Using the program LDHelmet, we estimated the population recombination rate *ρ* (= *2N_e_r*) for each of the nuclear chromosomes and the plastid. As expected, the plastid had much stronger LD and significantly lower *ρ* than observed in the nucleus (*ρ*_ptDNA_= 4.2 × 10^−4^, *ρ*_ntDNA_= 2.7 × 10^−3^, bootstrap test *p* <0.001). With our estimate of *N_e_* for the nuclear and plastid genomes, we can estimate the rates of recombination per cell division (r) from *r* = *ρ/*2*N_e_*. This calculation reveals that, in nature, the physical rate of recombination in the plastid is similar to the rate of recombination in the nuclear genome (Plastid *r* =8.2 × 10^−11^ /bp/generation, Nuclear *r*=9.2 × 10^−11^ /bp/generation, bootstrap test *p* = 0.64, Figure 2). Our finding seems unexpected, given the widely held view that the plastid is uniparentally inherited. However, there is clear evidence from genetic studies in *C. reinhardtii* that demonstrates nonreciprocal exchange between parental plastid genomes (e.g. Boynton et al. 1987; Lemieux and Lee 1987; Dürrenberger et al. 1996). If this form of gene conversion affects only short tracts of DNA it would explain why most of the plastid genome transmitted in any one cross is inherited from the mt+ parent. However, over evolutionary time, gene conversion can break down LD, especially between nearby loci.

Given that recombination rates of the plastid and nucleus appear to be similar, explaining the reduction in plastid *N_e_* by hitchhiking would require substantially more frequent or stronger selection in the plastid than the nucleus. One possible cause of increased hitchhiking in the plastid is the high gene density of the plastid genome, i.e., it contains 69 protein coding genes and 40 RNA genes, compared to an average nuclear region of the same size carrying only 28 protein coding genes. However, because nuclear genes are larger, the mean fraction of protein coding or RNA coding sites in the plastid genome is not substantially different from the average across the nuclear genome (plastid 43.3% vs. nuclear 35.5%). It is possible that the mutations in the plastid tend to have stronger selection coefficients causing either selective sweeps for beneficial changes (Maynard Smith and Haigh 1974) or background selection when mutations are deleterious (Charlesworth et al. 1993). Future investigation into the distribution of fitness effects of mutations across both genomes will be necessary to determine whether the reduction in *N_e_* in the plastid can be attributed to stronger effects of hitchhiking.

We have shown that rates of *de novo* mutation in the plastid and nuclear genomes are of similar magnitudes, in agreement with earlier studies based on DNA substitution patterns. This contrasts with the chloroplast genomes of land plants, where substitution rates are substantially lower than their nuclear counterparts. By integrating our direct, laboratory-based estimate of mutation with population genomic data, we have inferred that strong genetic drift, likely driven by hitchhiking, has reduced the amount of genetic diversity and increased linkage disequilibrium in the plastid genome. Our finding that the recombination rates of the plastid and nuclear genomes are similar means that in terms of population genetics we should not consider the plastid to be uniparentally inherited. This implies that the reduced diversity in the plastid genome is likely the results of stronger or more frequent selective sweeps and background selection. Increasing our knowledge of the role of mutation in shaping plastid genome evolution will require more comprehensive sampling among the millions of plastid-bearing species. As technical limitations associated with detecting *de novo* mutations are removed, opportunities will arise to investigate the spectrum and mechanisms of mutation in diverse plastid genomes and compare these to the nuclear genome.

---

**Supplemental Figure S1.**
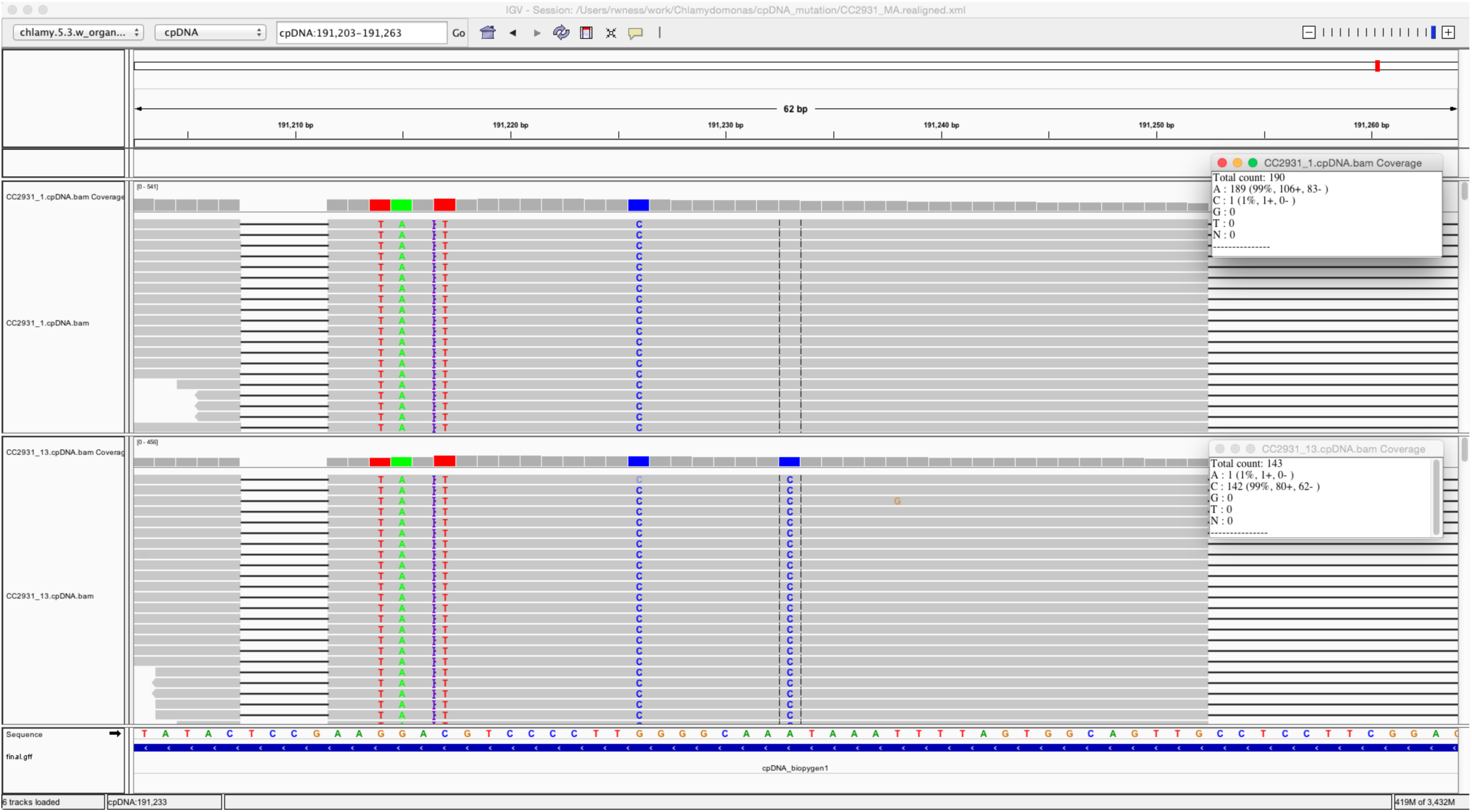
Screenshot from Integrated Genomics Viewer (IGV v2.3 Thorvaldsdóttir et al. 2012) of read alignment of mutation at position 191233 of the plastid genome. The bottom track is the sample that carries the *de novo* mutation (CC-2931 MA line 13) and the top track is a non-mutated MA line from the same ancestor (CC-2931 MA line 1). The putative A→C mutation is highlighted with vertical dashed lines in the centre of the image.

**Supplementary Table S1.** List of 20 natural isolates of *C. reinhardtii* sampled from two fields approximately 75km apart in Quebec, Canada. Strains were acquired from the *Chlamydomonas* collection (www.chlamycollection.org). The mating type of strains CC-3059 and CC-3062 are different from those reported in the Chlamydomonas collection based on sequence data from the *FUS1* and *MID* locus.

| Strain | Location | Mating type |
| --- | --- | --- |
| CC-3059 | Farnham, Quebec | mt- |
| CC-3060 | Farnham, Quebec | mt+ |
| CC-3061 | Farnham, Quebec | mt- |
| CC-3062 | Farnham, Quebec | mt- |
| CC-3063 | Farnham, Quebec | mt- |
| CC-3064 | Farnham, Quebec | mt+ |
| CC-3065 | Farnham, Quebec | mt+ |
| CC-3068 | Farnham, Quebec | mt+ |
| CC-3069 | Farnham, Quebec | mt+ |
| CC-3071 | Farnham, Quebec | mt+ |
| CC-3072 | Farnham, Quebec | mt+ |
| CC-3073 | Farnham, Quebec | mt- |
| CC-3075 | Macdonald campus, McGill University, Quebec | mt- |
| CC-3076 | Macdonald campus, McGill University, Quebec | mt+ |
| CC-3078 | Macdonald campus, McGill University, Quebec | mt+ |
| CC-3079 | Macdonald campus, McGill University, Quebec | mt- |
| CC-3082 | Macdonald campus, McGill University, Quebec | mt- |
| CC-3083 | Macdonald campus, McGill University, Quebec | mt- |
| CC-3084 | Macdonald campus, McGill University, Quebec | mt- |
| CC-3086 | Macdonald campus, McGill University, Quebec | mt+ |

## References

Aird D, Ross MG, Chen W-S, Danielsson M, Fennell T, Russ C, Jaffe DB, Nusbaum C, Gnirke A. 2011. Analyzing and minimizing PCR amplification bias in Illumina sequencing libraries. Genome Biology 12: R18.

Allen JF, Raven JA. 1996. Free-radical-induced mutation vs redox regulation: costs and benefits of genes in organelles. Journal of Molecular Evolution 42: 482–492.

Birky CW. 2001. The inheritance of genes in mitochondria and chloroplasts: laws, mechanisms, and models. Annual Review of Genetics 35: 125–148.

Boynton JE, Harris EH, Burkhart BD, Lamerson PM, Gillham NW. 1987. Transmission of mitochondrial and chloroplast genomes in crosses of *Chlamydomonas*. Proceedings of the National Academy of Sciences of the USA 84: 2391–2395.

Chamary JV, Parmley JL, Hurst LD. 2006. Hearing silence: non-neutral evolution at synonymous sites in mammals. Nature Reviews Genetics 7: 98–108.

Chan AH, Jenkins PA, Song YS. 2012. Genome-wide fine-scale recombination rate variation in Drosophila melanogaster. PLoS Genetics 8: e1003090.

Charlesworth B, Morgan MT, Charlesworth D. 1993. The effects of deleterious mutations on neutral molecular variation. Genetics 134: 1289–1303.

Denver DR, Morris K, Lynch M, Vassilieva LL, Thomas WK. 2000. High direct estimate of the mutation rate in the mitochondrial genome of Caenorhabditis elegans. Science (New York, NY) 289: 2342–2344.

Depristo MA, Banks E, Poplin R, et al. 2011. A framework for variation discovery and genotyping using next-generation DNA sequencing data. Nature Genetics 43: 491–498.

Dürrenberger F, Thompson AJ, Herrin DL, Rochaix JD. 1996. Double strand break-induced recombination in *Chlamydomonas reinhardtii* chloroplasts. Nucleic Acids Research 24: 3323–3331.

GuhaMajumdar M, Sears BB. 2005. Chloroplast DNA base substitutions: an experimental assessment. Molecular genetics and genomics 273: 177–183.

Haag-Liautard C, Coffey N, Houle D, Lynch M, Charlesworth B, Keightley PD. 2008. Direct Estimation of the Mitochondrial DNA Mutation Rate in Drosophila melanogaster. PLoS Biology 6: e204.

Howe DK, Baer CF, Denver DR. 2010. High Rate of Large Deletions in Caenorhabditis briggsae Mitochondrial Genome Mutation Processes. Genome Biology and Evolution 2: 29–38.

Howell N, Smejkal CB, Mackey DA, Chinnery PF, Turnbull DM, Herrnstadt C. 2003. The pedigree rate of sequence divergence in the human mitochondrial genome: there is a difference between phylogenetic and pedigree rates. American Journal of Human Genetics 72: 659–670.

Hua J, Smith DR, Borza T, Lee RW. 2012. Similar relative mutation rates in the three genetic compartments of Mesostigma and Chlamydomonas. Protist 163: 105–115.

Kelleher J, Ness RW, Halligan DL. 2013. Processing genome scale tabular data with Wormtable. BMC Bioinformatics 14: 356.

Kimura M. 1983. The neutral theory of molecular evolution. Cambridge, UK: Cambridge University Press.

Kong A, Frigge ML, Masson G, et al. 2012. Rate of *de novo* mutations and the importance of father’s age to disease risk. Nature 488: 471–475.

Lanfear R, Welch JJ, Bromham L. 2010. Watching the clock: Studying variation in rates of molecular evolution between species. Trends in Ecology & Evolution 25: 495–503.

Lemieux C, Lee RW. 1987. Nonreciprocal recombination between alleles of the chloroplast 23S rRNA gene in interspecific *Chlamydomonas* crosses. Proceedings of the National Academy of Sciences of the USA 84: 4166–4170.

Lesecque Y, Mouchiroud D, Duret L. 2013. GC-Biased Gene Conversion in Yeast Is Specifically Associated with Crossovers: Molecular Mechanisms and Evolutionary Significance. Molecular Biology and Evolution 30: 1409–1419.

Li H, Durbin R. 2009. Fast and accurate short read alignment with Burrows-Wheeler transform. Bioinformatics 25: 1754–1760.

Li H, Handsaker B, Wysoker A, Fennell T, Ruan J, Homer N, Marth G, Abecasis G, Durbin R. 2009. The sequence alignment/map format and SAMtools. Bioinformatics 25: 2078–2079.

Lynch M, Koskella B, Schaack S. 2006. Mutation pressure and the evolution of organelle genomic architecture. Science 311: 1727–1730.

Lynch M, Sung W, Morris K, et al. 2008. A genome-wide view of the spectrum of spontaneous mutations in yeast. Proceedings of the National Academy of Sciences of the USA 105: 9272–9277.

Maynard Smith J, Haigh J. 1974. The hitch-hiking effect of a favourable gene. Genetical Research 23: 23–25.

Mcvean GAT, Myers SR, Hunt S, Deloukas P, Bentley DR, Donnelly P. 2004. The fine-scale structure of recombination rate variation in the human genome. Science 304: 581–584.

Molina J, Hazzouri KM, Nickrent D, et al. 2014. Possible loss of the chloroplast genome in the parasitic flowering plant Rafflesia lagascae (Rafflesiaceae). Molecular Biology and Evolution 31: 793–803.

Morgan AD, Ness RW, Keightley PD, Colegrave N. 2014. Spontaneous mutation accumulation in multiple strains of the green alga, Chlamydomonas reinhardtii. Evolution 68: 2589–2602.

Ness RW, Morgan AD, Colegrave N, Keightley PD. 2012. An estimate of the spontaneous mutation rate in *Chlamydomonas reinhardtii*. Genetics 192: 1447–1454.

Ness RW, Morgan AD, Vasanthakrishnan RB, Colegrave N, Keightley PD. 2015. Extensive *de novo* mutation rate variation between individuals and across the genome of *Chlamydomonas reinhardtii*. Genome Research Advanced.

Race HL, Herrmann RG, Martin W. 1999. Why have organelles retained genomes? Trends in Genetics 15: 364–370.

Sloan DB, Taylor DR. 2011. Evolutionary Rate Variation in Organelle Genomes: The Role of Mutational Processes. In. Berlin, Heidelberg: Springer Berlin Heidelberg. p. 123–146.

Smith DR. 2015. Mutation rates in plastid genomes: they are lower than you might think. Genome Biology and Evolution.

Smith DR, Keeling PJ. 2012. Twenty-fold difference in evolutionary rates between the mitochondrial and plastid genomes of species with secondary red plastids. Journal of Eukaryotic Microbiology 59: 181–184.

Smith DR, Keeling PJ. 2015. Mitochondrial and plastid genome architecture: Reoccurring themes, but significant differences at the extremes. Proceedings of the National Academy of Sciences: 201422049.

Smith DR, Lee RW. 2014. A plastid without a genome: evidence from the nonphotosynthetic green algal genus Polytomella. Plant Physiology 164: 1812–1819.

Städler T, Haubold B, Merino C, Stephan W, Pfaffelhuber P. 2009. The impact of sampling schemes on the site frequency spectrum in nonequilibrium subdivided populations. Genetics 182: 205–216.

Sung W, Ackerman MS, Miller SF, Doak TG, Lynch M. 2012. Drift-barrier hypothesis and mutationrate evolution. Proceedings of the National Academy of Sciences of the USA 109: 18488–18492.

Thorvaldsdóttir H, Robinson JT, Mesirov JP. 2012. Integrative Genomics Viewer (IGV): highperformance genomics data visualization and exploration. Briefings in Bioinformatics.

Van Breusegem F, Dat JF. 2006. Reactive oxygen species in plant cell death. Plant Physiology 141: 384–390.

Vanwinkle-Swift KP. 1980. A model for the rapid vegetative segregation of multiple chloroplast genomes in *Chlamydomonas:* Assumptions and predictions of the model. Current Genetics 1: 113–125.

Zhu YO, Siegal ML, Hall DW, Petrov DA. 2014. Precise estimates of mutation rate and spectrum in yeast. Proceedings of the National Academy of Sciences of the USA 111: E2310–2318.

